# Machine-Learning Classification of Motor Unit Types in the Adult Mouse

**DOI:** 10.1101/2025.11.18.689075

**Authors:** María de Lourdes Martínez-Silva, Reuben M Ahorklo, Emily J Reedich, Rebecca D Imhoff-Manuel, Natallia Katenka, Marin Manuel

## Abstract

The diversity of motor units arises from differences in the contractile properties of muscle fibers and the intrinsic electrical properties of their motoneurons. In mice, however, this relationship has not been quantitatively defined, and conventional classification often relies on subjective thresholds. Here, we combined in vivo intracellular recordings with supervised and unsupervised machine-learning methods to test whether motoneuron electrophysiology can predict the physiological identity of mouse motor units. Unbiased clustering identified four groups corresponding to slow (S), fast fatigue-resistant (FR), intermediate (FI), and fast fatigable (FF) types. A multinomial logistic regression model performed well, with most errors occurring between FI and FF types, which showed substantial overlap. Reducing the task to three classes improved accuracy. Feature selection revealed that four electrophysiological properties (input conductance, rheobase, AHP duration, maximal frequency) were sufficient for high predictive performance. Overall, this study provides a quantitative description of mouse motor-unit properties and a framework for incorporating motor-unit diversity into future investigations of neuromuscular physiology and disease.

## Introduction

Motor units (the motoneuron together with all the muscle fibers it innervates) form the fundamental building blocks of motor control [1]. The functional diversity of motor units underlies the ability of muscles to perform both fine, slow, deliberate tasks and powerful, rapid movements [2]. In vertebrates, motor units were classically grouped into at least three types: slow (S), fast fatigue-resistant (FR), and fast fatigable (FF) types, which differ fundamentally in contractile speed, force generation, and fatigue behavior [3]. Seminal work by Burke and colleagues laid the foundation for this classification by correlating mechanical responses, such as twitch contraction time, force, and fatigue indices, with distinct histochemical properties [2], [3]. Slow units are characterized by prolonged contraction, high oxidative capacity, and strong resistance to fatigue, crucial for posture and endurance. Fast fatigable units provide high force production and rapid contraction but are notably susceptible to fatigue due to their reliance on glycolytic metabolism [1]. However, some units were found to not fit exactly into these categories and a fourth intermediate class (FI), was proposed [4], [5], [6], [7]. The FI classification acknowledges the gradation of fatigue resistance that could not be adequately described by the simple FR/FF dichotomy [5], [8], [9], [10], [11].

Concurrently, on the neural side, motoneurons (MNs) exhibit electrical properties that correlate systematically with the contractile characteristics of their motor units [3]. Thanks to pioneering work in feline triceps motoneurons, and subsequent studies in rats, it is well established that motoneuron input resistance, afterhyperpolarization (AHP) duration, axonal conduction velocity, and recruitment thresholds differ systematically across motor unit types (S, FR, FF) and correlate with twitch contraction time, force, and fatigue profile [12], [13], [14]. Generally, S motoneurons are more excitable, having high input resistance and lower rheobase, while fast units (FF and FR) have lower input resistance and higher rheobase (see [13] for a recent review). This systematic relationship underpins fundamental concepts such as Henneman’s size principle governing orderly recruitment [1].

More recently, we were able to perform recordings in adult mice *in vivo* and study the contractile properties of Triceps Surae (TS) motor units together with measurements of motoneuron electrical properties in two different mouse models of Amyotrophic Lateral Sclerosis. In this work, we classified the recorded motoneurons in the three classical types (S, FR and FF) using hand-picked thresholds in twitch contraction time and twitch amplitude [15]. This work was the first of its kind performed in mice, and opens the door to studying the properties of mouse motor units in healthy mice, but also in mouse models of various human diseases.

However, several gaps remain. First, although the mouse has become a crucial model for the study of human disorders, their motor units display faster contraction kinetics and higher maximal firing frequencies compared to rats and cats, illustrating that it is critical to fully characterize the properties of motor units and their motoneurons in this species [15], [16]. Second, while threshold-based classification (e.g. using twitch contraction times or fatigue indices) is common, the inherent overlap and continuous nature of motor unit properties means that classification schemes are subject to ambiguity [4], [7]. Third, with advances in computational methods, machine learning offers an opportunity to build robust predictive models.

In this paper, we address these gaps by measuring a large panel of contractile properties of individual motor units in the mouse Triceps Surae, classifying them into physiological types based on their contractile properties using unbiased, unsupervised clustering algorithms, while recording a comprehensive set of motoneuron electrical properties. The main One goal of this study was then to fit a statistical model allowing the prediction of a motor unit type based on the electrical signature of its motoneuron. This approach deepens our understanding of neuromuscular organization and enables faster, less invasive classification in mice and mouse models of neuromuscular disorders.

## Methods

### Animals

All experiments done in Paris were performed in accordance with European directives (86/609/ CEE and 2010-63-UE) and the French legislation. They were approved by Paris Descartes University ethics committee (authorization CEEA34.MM.064.12). Experiments done at the University of Rhode Island were performed according to guidelines of the National Institutes of Health guide for the care and use of Laboratory animals (NIH Publications No. 8023, revised 1978) and have been authorized by the University of Rhode Island’s Animal Care and Use Committee (Protocol AN2021-018).

27 B6SJL mice (age 60±5.2 d [48–71 d] N=27; weight 25±3.6 g [17–32 g] N=27; 17 males and 10 females) were used for this study. Mice were purchased from the Jackson Laboratory and bred in house with a 12 h/12 h light/dark cycle and food and water *ad libitum*.

### Surgical procedure

The surgical procedure has been described previously [15], [17]. Briefly, atropine (0.25 mg/ kg; Covetrus) and methylprednisolone (2 mg/kg; Covetrus) were given subcutaneously to prevent salivation and edema, respectively. Anesthesia was induced with an i.p. injection of Fentanyl 0.025 mg/kg (Covetrus), Midazolam 7.5 mg/kg (Covetrus) and Medetomidine 0.5 mg/ kg (Covetrus). The heart rate was monitored with an EKG, and the central temperature was kept around 37°C using an infrared heating lamp and an electric blanket. Then, the mouse was artificially ventilated with pure oxygen (SAR-1000 ventilator; CWE) through a cannula inserted in the trachea. The ventilator settings were adjusted to maintain the end-tidal CO_2_ level at ∼4 % (Micro-Capstar; CWE). One catheter was introduced in each of the left and right external jugular veins. The first one was used to deliver supplemental doses of anesthesia whenever necessary (usually every 20–30 min) by i.v. injection (10 % of the dose used for anesthesia induction). The adequacy of anesthesia was assessed by lack of noxious reflexes and stability of the heart rate (usually 400–500 bpm) and end-tidal PCO_2_. The other catheter was used for a continuous perfusion of saline (100 µL/h, Covetrus). The vertebral columns were immobilized with two pairs of horizontal bars (Cunningham Spinal Adaptor; Stoelting, Dublin, Ireland) applied on the Th12 and L2 vertebral bodies, and the L2–L4 spinal segments were exposed by a laminectomy at the T13–L1 level. A custom made chamber was fit around the exposed spinal segments and silicon sealant (Body Double Fast Set, Smooth-On Inc.) was applied to create a recording chamber. This chamber was filled with mineral oil to prevent desiccation of the spinal cord. All branches of the sciatic nerve were cut except for the nerve innervating the Triceps Surae (TS), which was left intact and mounted on a mono-polar stimulation electrode. The tendon of the TS was dissected free and securely attached to an isometric force transducer (Aurora 404C, Aurora Scientific). The skin of the leg was pulled taught and used to form a pool filled with mineral oil around the muscle. The temperature of the muscle was maintained around 37 °C (36±1 °C [33-40 °C] N=131) using a recirculating flow of mineral oil heated with an inline heater (SF-28, Warner Instruments) and a temperature probe inserted underneath the muscle.

Intracellular recordings were performed using glass microelectrodes (tip diameter 1.0–1.5 µm) filled with KCl 3 M (resistance 8–15 MΩ). Intracellular recordings were obtained with an Axoclamp 2B amplifier and Spike2 software (CED, Cambridge, England). All recordings were obtained using discontinuous current clamp (7–9 kHz [18]) and sampled at 30 kHz. At the beginning of the experiment, the initial length of the muscle was set to the length at which the ankle was flexed at 90°. The length was then adjusted in order to record the maximal muscle twitch amplitude. Electromyographic activity (i.e. single motor unit compound action potential, MUAP) was recorded at the same time using fine stainless steel wires inserted underneath the muscle fascia (signal amplified 100–1000x, and band-pass filtered at 10 Hz–10 kHz; AM System Model 1700).

### Electrophysiological recordings

Upon impalement of a motoneuron, we identified the motoneuron as innervating TS muscle fibers based on the presence of an antidromic spike in response to the stimulation of the nerve and its ability to induce muscle twitches upon injection of current through the microelectrode. A series of tests was conducted to characterize the contractile properties of the motor unit and the electrophysiological properties of the motoneuron. First, a series of 10, 1 ms long, pulses of current (repeated at 1 Hz) were used to elicit single action potentials and produce individual twitches of the motor unit. Then, 500 ms-long trains of 1 ms-long current pulses at 20, 25, 30, 35, 40, 50, 60, 70, 80, and 150 Hz (separated by 30 s pauses) were used to study the unfused and fused tetanus of the motor unit. The intensity of the pulses was adjusted by hand in order to reliably elicit one action potential on each pulse. Trains where some action potentials failed to be triggered, or where the the MUAPs were not constant were excluded from analysis. Finally, a new series of 1 ms pulses at 1 Hz were used to elicit single twitches and quantify the amount of post-tetanic potentiation. The input resistance/conductance of the motoneuron was measured from the peak and plateau responses generated by small-amplitude 500 ms square current pulses [19]. The membrane time constant was measured on the relaxation of the membrane potential following 1 ms hyperpolarizing current pulses using the “peeling” method [20]. The motoneurons’ firing characteristics were examined using triangular current ramps applied at a rate of 0.1–2 nA/s. The total ramp duration depended on the ramp speed and the current intensity reached at the top of the ramp, which was adjusted by hand based on the rheobase of the motoneuron. The instantaneous firing frequency was plotted against the injected current intensity at the moment of the spike. The voltage threshold for firing was measured on the first spike of the ramp, and was defined as the point where the first derivative of the voltage reached 10 mV/ms [21]; the recruitment current, corresponding to the intensity of current at the voltage threshold; the de-recruitment current, as the current intensity at the time of the last spike; the ΔI values, corresponding to the difference between de-recruitment and recruitment currents; the frequency of the first inter-spike interval (onset firing frequency); the maximal frequency reached during the ramp; the firing frequency of the last inter-spike interval (offset firing frequency); the ΔF value, corresponding to the difference between the offset and onset frequencies; the gain, calculated as the slope of a linear regression applied over the “primary range” of the *f-I* curve [19]. The rheobase was measured in response to 20 ms-long pulses of current and was defined as the intensity of current necessary to elicit an action potential 50 % of the time. 5 s-long current pulses at intensities close to the rheobase were used to determine if the firing of the motoneuron was delayed or immediate [22]. Finally, trains of eight pulses of current at 40 Hz, repeated every second, were delivered for 3 minutes (180 trains in total) in order to test the fatigability of the motor unit. As shown previously [15], in some motor units the MUAPs had a tendency to decrease over the course of a single train. Therefore, we measured the amplitude of the first twitch of each train, and the fatigue index was calculated as the ratio of the amplitudes of the first twitch of the first train to the last train [15].

### Data Selection

Only motoneurons with a membrane potential more hyperpolarized than −50 mV, an overshooting action potential, and a membrane time constant > 1.5 ms were kept for analysis. Overall, our final population consisted of 131 motor units. No data were excluded as outliers, in order to preserve the full range of biological variability. The number of tests that could be completed in each cell was variable, however. For the clustering analysis (see below), we first used a sub-sample of 78 motor units in which the full range of contractile properties were recorded, before extending the classification to the remainder of the units. For building the statistical classification model (see below), we focused on another sub-sample of 103 motoneurons (out of 131), in which the most important electrical properties were fully characterized (see below).

### Motor unit clustering

To group the motor units in one of the four types (S, FR, FI and FR), we used principal component analysis (PCA) to reduce the dimensionality of the dataset (78 observations with 36 features). First the features were standardized (`StandardScaler`) and then the first five principal compo-nents were computed (`PCA(n=5)`). We then used K-means clustering to group the recordings in one of 4 clusters (`KMeans(n=4)`). To classify the remainder of the motor units (53 motor units in which some of the features were missing), we used the features available for each of the remaining units to train a k-nearest-neighbor classifier in order to find the cluster to which each remaining unit was the closest. We used a value of 𝑘 = 3 with a Cartesian distance weighting to take into account more variability than simply the nearest observation.

### Statistical analysis

Based on the procedure above, each unit could be classified in one of four classes, and we then compared the average properties of the units in each cluster vs. the other clusters. Comparisons between the four clusters were performed on the basis of the effect size of the difference of mean and its 95 % confidence interval. Since some of the features were not following a normal distribution, we performed the non-parametric Kruskal-Wallis H-test for independent samples. If this test reveals that the four groups were different, then a pairwise post-hoc Mann-Whitney U-test with Holme’s adjustment for family-wise error rate was performed. Linear relationships were quantified using a linear least-squares regression, and we reported the value of the coefficient of determination 𝑟^2^ and the p-value for a hypothesis test whose null hypothesis was that the slope was zero, using Wald Test. All values are reported as mean ± standard deviation. A summary of each property is provided in Supplemental Table 1 and Supplemental Table 2. For each cluster, we provided the mean ± standard deviation, the range of the values and the number of observations. We also provided the H value and *p* value of a Kruskal-Wallis test of the equivalence of the means. We then performed pairwise comparisons between each cluster. For each pair, we showed the effect size (Hedges’ g, or the mean difference if hedges’ g could not be calculated) and the 95 % confidence interval around the effect size. If the Kruskal-Wallis test showed a difference between groups, we performed a Mann-Whitney test and provided the U value and p value (adjusted using Holmes’ method to control the family-wise error rate).

Statistical analysis was performed in Python (v3.12), NumPy (v2.2 [23]), Pandas (v2.3 [24]), Seaborn (v0.13 [25]), matplotlib (v3.10 [26]), Pingouin (v0.5 [27]), dabest (v2025.03.27 [28]), scipy (v1.16 [29]).

### Motor unit type prediction

#### Feature selection and preprocessing

To predict motor unit type (S, FR, FI, and FF) from intrinsic motoneuron properties, we selected a set of 23 electrophysiological variables that are commonly recorded during intracellular recordings and were consistently available across the dataset. These features included resting membrane potential, input resistance (measured at the peak and plateau [19], [30]), sag ratio, input conductance (calculated as the inverse of the peak input resistance), membrane time constant, current at firing onset, current at firing offset, ΔI, onset firing frequency, ramp maximum frequency, offset firing frequency, ΔF, f-I curve gain (in the primary range), voltage threshold (absolute value), voltage threshold (relative to resting membrane potential), *f-I* curve profile (clockwise, anti-clockwise, or symmetrical), axonal conduction velocity, rheobase, spike height, spike width, AHP amplitude, and AHP half relaxation time. We were able to measure these 23 properties in 103 motoneurons out of the total population of 131 units, and we used that sub-sample to train our predictive algorithms.

#### Model training and hyperparameter optimization

To test whether electrophysiological properties could predict motor unit type, multiple supervised learning classification algorithms were trained on the 103 observations, each with 23 features. The models we evaluated were multinomial logistic regression (`LogisticRegression` in scikit-learn), random forest (`RandomForestClassifier`), gradient boosting (`GradientBoostingClassifier`), XGBoost (`XGBClassifier`), support vector machine (`SVC`), and k-nearest neighbors (`KNeighborsClassifier`). Each algorithm was implemented within a scikit-learn (v.1.7.2) pipeline that involved standardization of the numerical features and one-hot encoding of the categorical features. Model performance was evaluated using five-fold stratified cross-validation, maintaining the relative distribution of motor unit types across folds.

For each model, we performed a systematic grid search to optimize their hyperparameters using the `GridSearchCV` function from scikit-learn. Each grid search used balanced accuracy as the primary scoring metric, as it compensates for unequal class frequencies among the four motor unit types. The macro F1-score was also computed to assess the model’s performance consistency across classes. To address class imbalance, all models that supported it (multinomial logistic regression, SVM, random forest) used class-weight balancing. The final model was refitted using the best hyperparameters determined by cross-validation.

The final model that was retained for further analyzes was a multinomial logistic regression model because of its performance and its interpretability. Odds ratios were calculated as the exponent of the coefficients obtained from the fitted model. To estimate the uncertainty of the coefficients, we performed bootstrapping by resampling the dataset with replacement (1000 iterations), refitting the complete model for each bootstrap sample, and computing the empirical 2.5th and 97.5th percentiles of the coefficient distributions as 95% confidence intervals. Confidence intervals were then exponentiated to obtain the corresponding intervals for the odds ratios.

Confusion matrices were generated to visualize class-level performance, particularly for distinguishing between the four motor unit types (FF, FI, FR, S). An additional three-class variant of the problem—grouping FF and FI together (fast), and treating FR and S as distinct classes—was evaluated to examine whether coarser classification improved separability.

All analyses were conducted in Python (v3.12) using scikit-learn (v1.7 [31]), NumPy (v2.2 [23]), Pandas (v2.3 [24]), seaborn (v0.13 [25]), and matplotlib (v3.10 [26]).

## Results

### Motor unit classification

We recorded a total of 131 motor units *in vivo* from the Triceps Surae (TS) muscle of B6SJL mice and characterized both their contractile and electrophysiological properties using an anesthetized but non-paralyzed adult mouse preparation [17], [32]. Instead of hand-picked thresholds as we did previously [15], we decided to apply unsupervised clustering algorithms to group motor units that behaved similarly. We used 36 contractile variables, including twitch amplitude, contraction and half-relaxation times, tetanic force, fatigue index, force-frequency curves and many others. Unfortunately, due to the inherent instability associated with performing these recordings *in vivo*, it was not always possible to record all 36 parameters in all recorded motor units. We therefore focused classification on a subset of 78 units in which these 36 parameters had been recorded.

We first performed principal component analysis (PCA) on the standardized contractile variables. The first five principal components explained 80.6 % of the total variance (PC1=48.9 %, PC2=15.0 %, PC3=15.0 %, PC4=15.0 %, PC5=3.9 %), indicating that the dimensionality of the dataset could be effectively reduced to a few components (Figure 1D2). PC1 and PC2 primarily reflected differences in contraction speed and force production. PC1, which captured almost half of the variance, correlated positively with twitch and tetanic force, whereas PC2 captured differences in contraction and relaxation kinetics (Figure 1D3. See also Supplemental Table 3 for a complete list) Based on the classical grouping of motor units in S, FR, FI and FF types [4], we used K-means clustering (𝑘 = 4) to generate four distinct groups of motor units (Figure 1D1). The clusters were well separated in the PCA space. To assign the remaining partially characterized units (N=53) to one of these four clusters, we trained a k-nearest-neighbor classifier (𝑘 = 3, distance-weighted) on the complete subset and used it to predict the most likely class of each unclassified observation.

**Figure 1.**
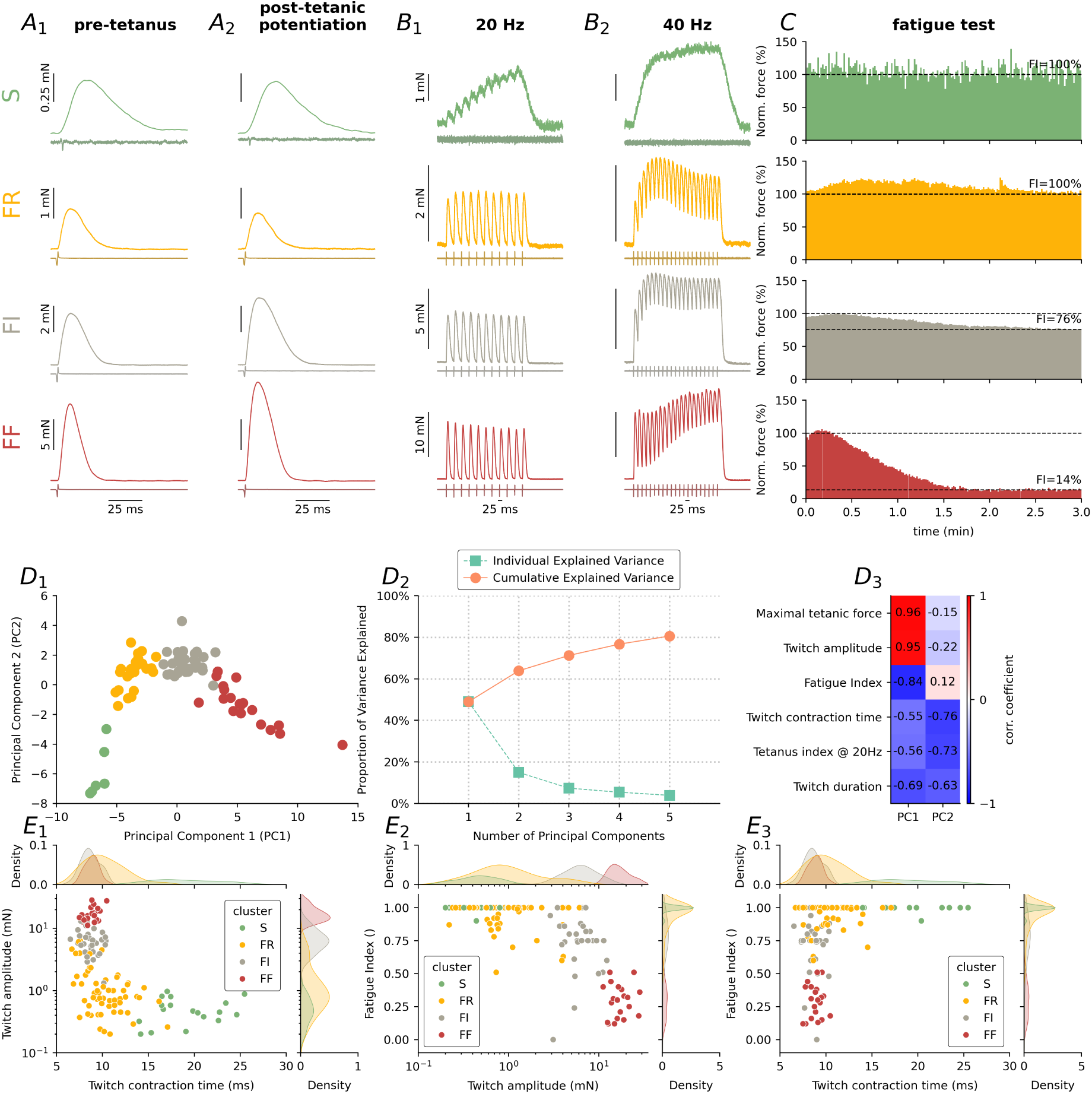
Classification of motor units in four groups using unsupervised clustering algorithm. **A** Representative examples of force recordings elicited by injecting short pulses of current in the cell body of the motoneurons in the four clusters of motor units. In each sub-panel, the top trace is the force, and the bottom lighter trace is the MUAP recording. **A_1_** show single twitches that were obtained just after the impalement of the motoneurons, while **A_2_** are traces recorded after maximal contraction of the motor unit was elicited by 150 Hz stimulation of the motoneuron. Scale bars are the same in A_1_ and A_2_. Traces are averages of 5-10 sweeps. **B** Examples of response of the same motor units in A to 20 Hz (**B_1_**) or 40 Hz (**B_2_**) stimulations, showing the 3 types of sag observed in those units. Scale bars are the same in B_1_ and B_2_. **C** Evolution of the force generated by the motor units in response to 8 pulses of current at 40 Hz, repeated every 1 sec for 3 min. **D_1_** Scatter plot of the first two principal components. Colors correspond to the clusters identified by K-means. **D_2_** Scree plot showing the variance explained by each principal component. **D_3_** Loading values of the first two principal components for a few representative contractile parameters. A complete list of loadings is available at Supplemental Table 3. **E_1_** Distribution of the twitch amplitude (logarithmic scale) vs. twitch contraction time. Marginal plots on the top and right side show the density distribution of each cluster, while the central scatter plot shows the individual motor units, color coded by cluster. **E_2_** Distribution of the Fatigue Index vs. twitch amplitude (logarithmic scale). Same organization as in D_1_. **E_3_** Distribution of the Fatigue Index vs. twitch contraction time. Same organization as in D_1_.

### Contractile properties of the identified mouse motor unit types

Our strategy grouped motor units into clusters that matched the expected properties of the different types of motor units described in cats [3], rats [33], [34], [35], and mice [15]: One cluster had small twitch amplitudes, contracted slowly, and showed strong fatigue resistance, probably corresponding to type S motor units (Figure 1A-C, N=18 units, 14 % of the population). On the other hand, another cluster contained motor units with the strongest twitches, a short contraction time, and high fatigability, which corresponds to the properties of type FF motor units (Figure 1A-C, N=21 units, 16 %). A third cluster had small to medium twitch amplitudes, contraction times that were faster than the S group, but more spread than the FF group, and were mostly resistant to fatigue, which fits well with the type FR motor units (Figure 1A-C, N=59 units, 45 %). Finally, the fourth cluster had medium to high twitch amplitudes, fast contraction times, and intermediate fatigue resistance, which can be classified as an intermediate type FI, between FR and FF units (Figure 1A-C, N=33 units, 25 %) (Figure 1E_1-3_). The distribution of the different types matches the proportions we had observed previously [15].

Other properties also matched the expected distribution among the different types (see Figure 2 for the most relevant. A full comparison of all the properties is available in Supplemental Table 1 and Supplemental Figure 2): maximal tetanic force (Figure 2E) was smallest in S, followed by FR, then FI and FF had the largest. Yet, the ratio between twitch amplitude and tetanic force (Figure 2F) was the largest in S and FR units, and smallest in FI and FF units, indicating that the summation of contractions during high-frequency stimulation was more efficient in the slowest units (see also Figure 3). MUAP amplitudes (Figure 2G) were also distributed in the expected order (S ≈ FR > FI > FF). Axonal conduction velocity (Figure 2H) was slowest in the S units, followed by FR and FI, but was not different between FI and FF. Post-tetanic potentiation (Figure 2I) was assessed based on the twitch amplitude recorded after a series of unfused and fused tetanic contractions. FF and FI units were potentiated by an average of 129 % and 128 %, respectively, while S and FR units did not potentiate on average (although there was a correlation between potentiation and twitch amplitude 𝑟^2^ = 0.40, p = 9.5⋅10−11 such that the largest FR units tended to potentiate more than the smallest).

**Figure 2.**
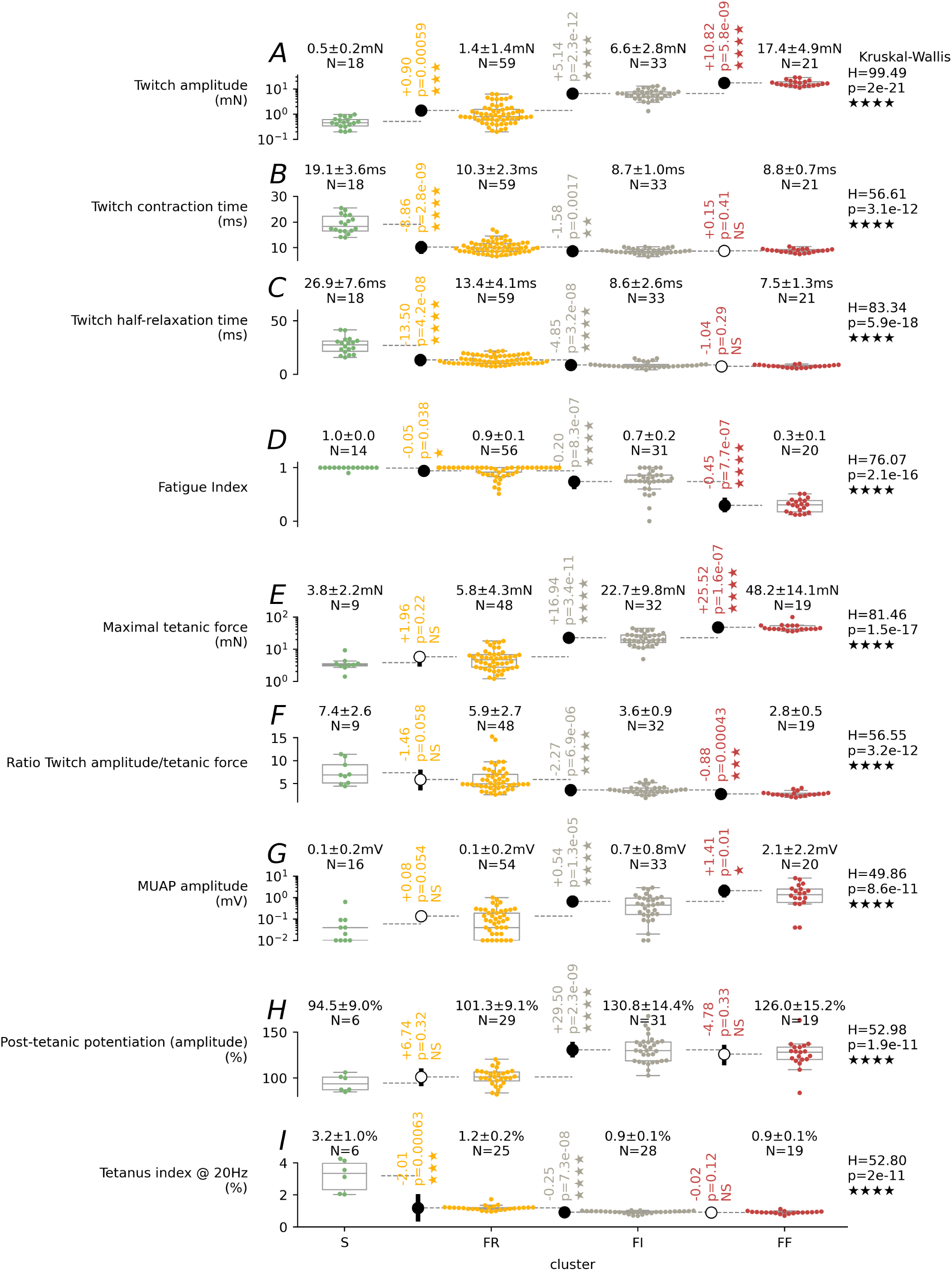
Distribution of the major contractile properties across the different types. **A-I** For each property (row) and cluster (columns), we show the individual data points superimposed over a boxplot that shows the three quartile values of the distribution along with extreme values. The “whiskers” extend to points that lie within 1.5 x inter-quartile distance of the lower and upper quartile. Above each boxplot, we report the mean ± standard deviation of each of the cluster sub-population. On the far right of the plot is the result of the Kruskal-Wallis H-test for independent samples, showing the H statistic and the *p* value. In between each cluster, we show the difference of the means (shown with dashed lines) between the group to the left and the right, with its associated 95 % confidence interval. The *p* value corresponds to the result of the pairwise Mann-Whitney U Test post-hoc test with Holm adjustment for family-wise error rate. If the Kruskal-Wallis test shows that the four populations have different means, and the Mann-Whitney test is statistically significant, then the difference of means is shown with a filled symbol, otherwise, it is shown with an empty symbol. *p* < 0.05: ★, *p* < 0.01: ★★, *p* < 0.001: ★★★, *p* < 0.0001: ★★★★.

**Figure 3.**
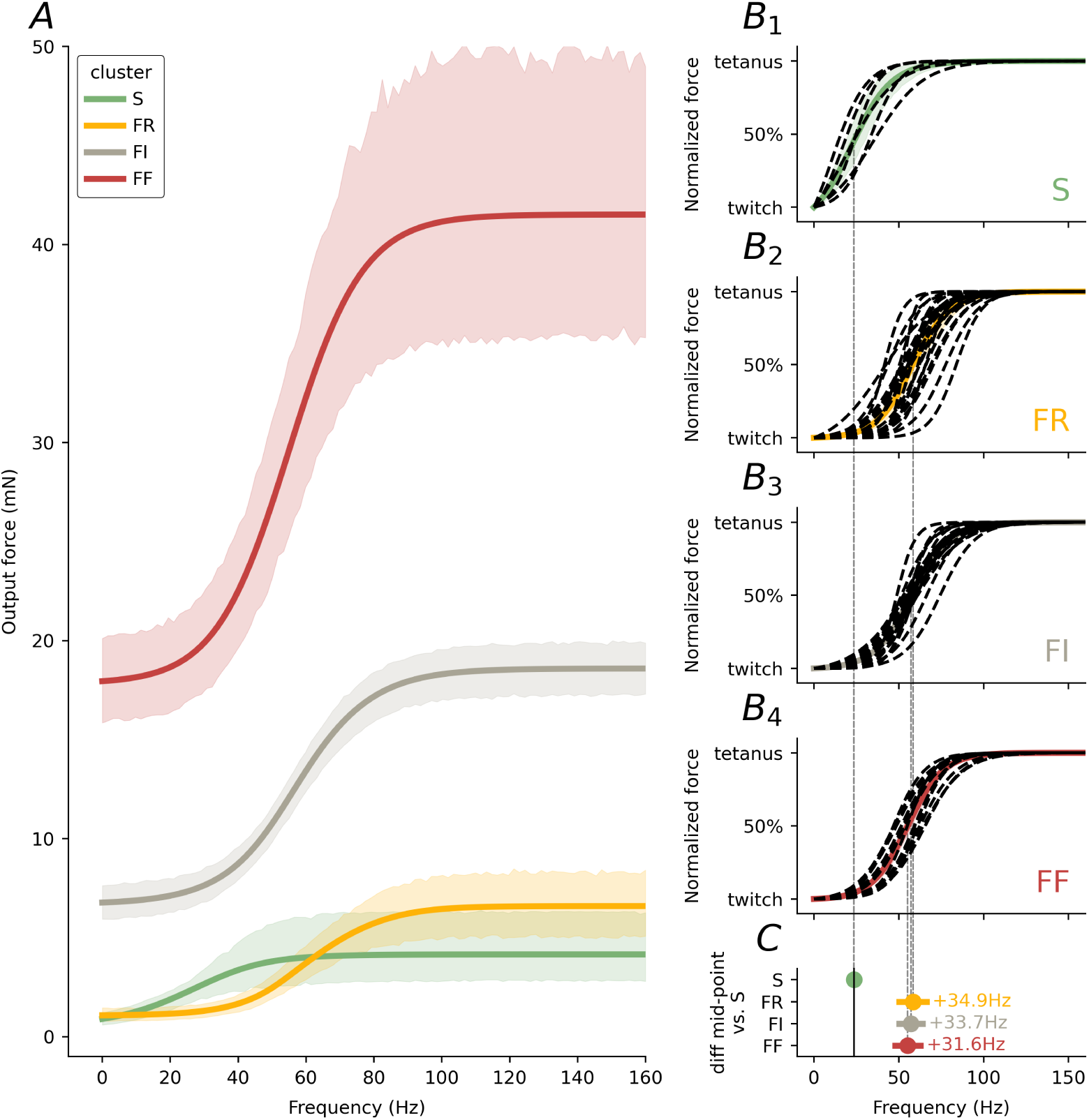
Force-Frequency curves of the motor units. **A** Non-normalized average curves for each of the four clusters of motor units. **B_1-4_** Force-Freq curves normalized by the minimum and maximum force. The dashed lines show the curves for individual motor units, while the thick line is the average curve. **C** Comparison of the mid-point frequency of each of the average curves with the S curve. Each point shows the mean difference with its 95 % confidence interval.

We studied the stimulation frequency *vs.* output force relationship of our motor units over the range 20–150 Hz (Figure 3). As expected based on their long contraction time and relaxation times (Figure 2), the force of S units started to sum at lower frequency and reached peak force at lower frequency as well. The force-frequency curve of the FR units started at the same force level, but started increasing at higher frequencies (Figure 3C) and reached a higher force output. Although, on average, the frequency at the curve mid-point is not different between the FR, FI and FF units (Figure 3C), observation of the individual curves show that a number of FR units start increasing at lower frequencies than FI and FF units (compare Figure 3B2 and B_3-4_), as expected based on the larger range of contraction times and relaxation times in FR units (Figure 2B-C).

During unfused tetanic contractions, the force often exhibited a “sag”: tetanic tension increased rapidly at the onset of the train, followed by a decrease of tension to reach a lower steady state, particularly in F type motor units [36]. We observed that the sag phenomenon, like in other species, was frequency dependent and occurred in three distinct forms (Figure 1B_1-2_): either the force profile did not show any sag at all, or it sagged to a steady state level (“simple” spike), or, after an initial dip, the force started to increase again (“complex” sag [37], [38]). As shown in Figure 4, sag was mainly observed in fast units, but not in all fast units. In response to a train of action potential at 20 Hz, none of the S motor units exhibited sag, while sag was starting to be apparent in some FR units. Note that 20 % is too low a frequency to allow fusion of twitches in the fastest units, and their sag profiles could therefore not be studied. As the train frequency increases, the proportion of units exhibiting sag increases, and more and more units exhibit complex sags, particularly in the FF and FI types (Figure 4B). In studies in rats, it was suggested that the ratio between the last and first twitch amplitude of the train (“tetanus index” [39]) could be used to identify S from F units. In the mouse, we also found that the tetanus index at 20 Hz and 30 Hz could be used to separate S from F units, although the separation was not very large between the fastest of the S units and the slowest of the FR units.

**Figure 4.**
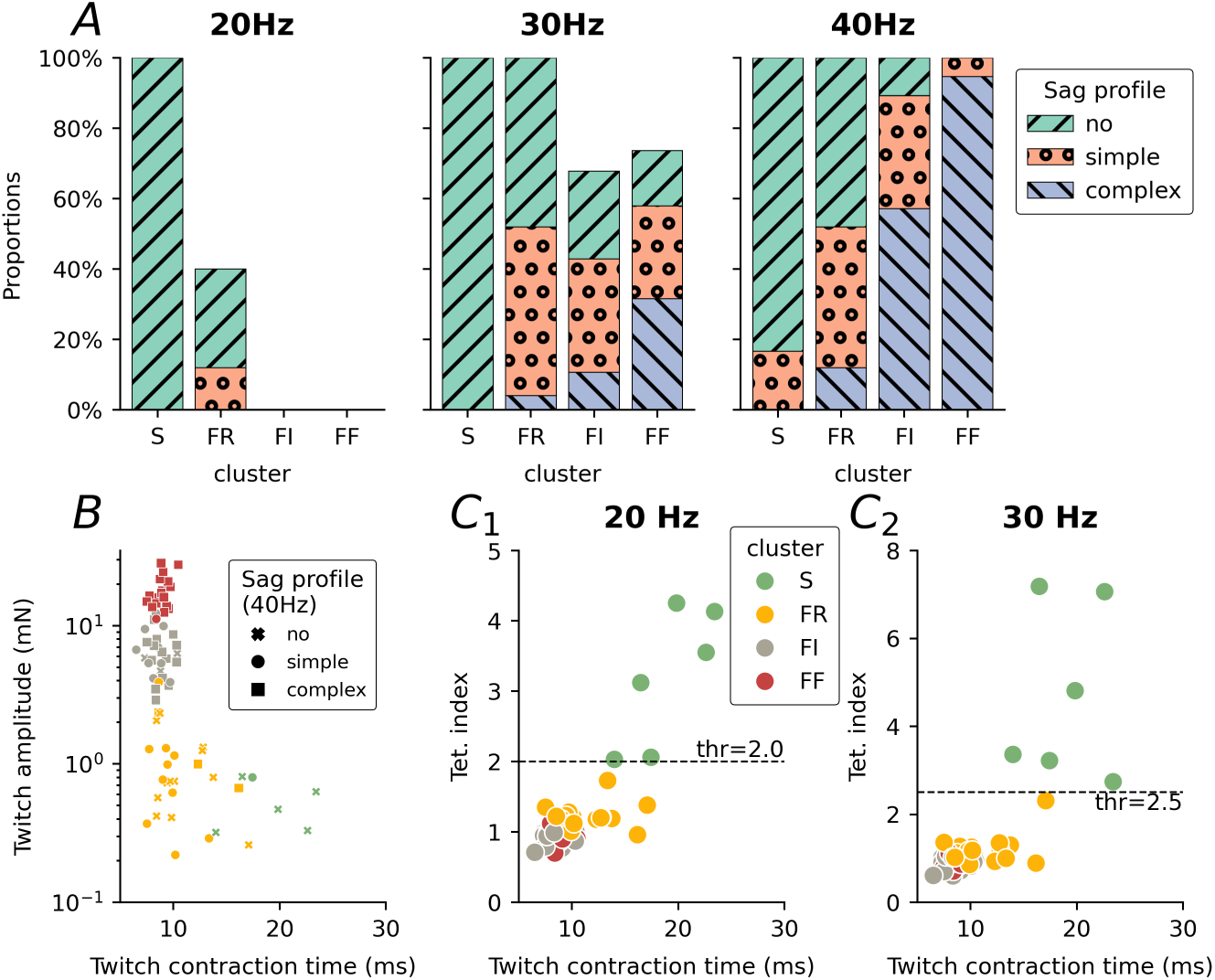
Sag profile of mouse motor units. **A** Proportion of the three types of sag profiles (see Figure 1B_1-2_ for examples) in each of the four types of motor units, as a function of the frequency of the stimulation. Only trains in which fusion of the twitches was observed were kept for analysis. **B** Distribution of the different sag profiles as a function of the twitch amplitude and twitch contraction time. **C** Tetanus index (ratio of last twitch over first twitch of the train) as a function of the twitch contraction time, measured at 20 Hz (**C_1_**) and 30 Hz (**C_2_**). The horizontal dashed lines represents a value that separates the S-type motor units from the F-type.

We wanted to test whether animal sex and/or muscle size affects our classification. For the effect of sex, we separated our population based on sex, and tested whether the motor unit forces were different between males and females (Supplemental Figure 1). The only difference we observed is that female FF units tended to exhibit smaller twitch amplitudes compared to males (hedge’s *g* = −1.94 [95 %CI −3.0–−0.88], p = 0.002, Supplemental Figure 1), but there were no differences in twitch contraction time, or tetanic force. A mixed-effects model including sex, body weight, and their interaction as fixed effects, and animal as a random intercept (Supplemental Figure 1), revealed no significant effects on twitch amplitude. Neither sex (β = 6.42 ± 13.40, p = 0.63) nor body weight (β = 0.03 ± 0.36, p = 0.94) significantly predicted twitch amplitude, and the interaction between sex and body weight was not significant (β = – 0.18 ± 0.53, p = 0.74). This observation suggests that, although differences in total force output are to be expected between males and females [40], [41], [42], these differences are minimal at the single motor-unit level, and are only observable in the largest FF motor units.

**Supplemental Figure 1.**
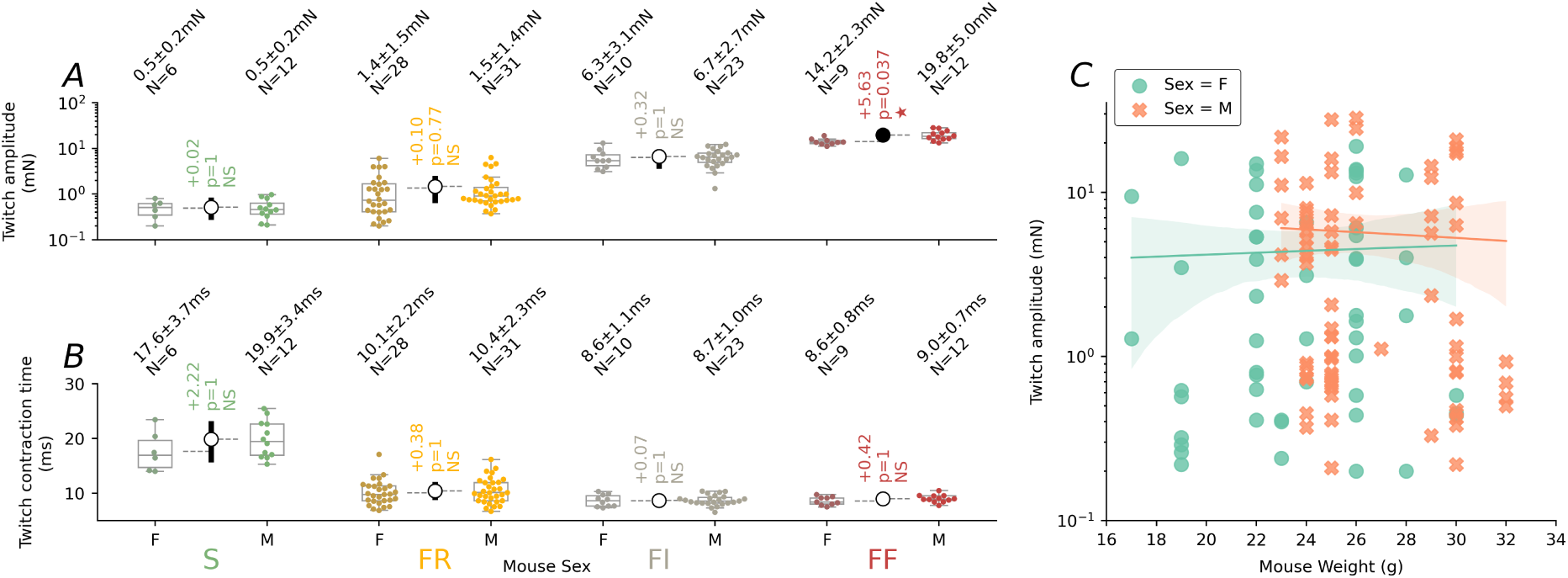
Effect of sex and mouse weight on motor unit force. **A** Twitch amplitude in each cluster, separated by sex. For each group, we show the individual data points superimposed over a boxplot that shows the three quartile values of the distribution along with extreme values. The “whiskers” extend to points that lie within 1.5 x inter-quartile distance of the lower and upper quartile. Above each boxplot, we report the mean ± standard deviation of each of the cluster sub-population. In between each sex, we show the difference of the means (shown with dashed lines) between the group to the left and the right, with its associated 95 % confidence interval. The *p* value corresponds to the result of the pairwise Mann-Whitney U Test post-hoc test with Holm adjustment for family-wise error rate. If the Mann-Whitney test is statistically significant, then the difference of means is shown with a filled symbol, otherwise, it is shown with an empty symbol. **B** Twitch contraction time in each cluster, separated by sex. Same organization as in A. **C** Scatter plot of the twitch amplitudes as a function of mouse weight (assumed to be proportional to the muscle weight), separated by sex. The regression lines are almost horizontal and largely overlapping, indicating an absence of effect of sex and body weight on the twitch amplitude.

### Electrophysiological properties of type-identified motoneurons

We next examined the electrophysiological properties of the motoneurons innervating each of these motor units. As expected, there were strong correlations between the electrical properties of the motoneurons and their physiological types (Figure 5). The passive properties of the motoneurons were different between the types of motor units [13]. S-type motoneurons had the largest input resistance, consistent with their smaller size [43], and a distinct absence of sag in their response to subthreshold current pulses [12], [44]. On the opposite end of the spectrum, FF-type motoneurons had the lowest input resistance, and their voltage responses displayed a pronounced sag (Figure 5A, Figure 5C–E). The rheobase (the minimum amount of current necessary to elicit an action potential) was directly related to the input resistance (𝑟^2^ = 0.57, p = 1.6⋅10−20): S motoneurons had the lowest rheobase, followed by FR, then FI and FF, which had similar values (Figure 5G). The after-hyperpolarization (AHP) following each spike is also known to be different between types [12], [44]. Unexpectedly, we did not observe any difference in the amplitudes of the AHP (Figure 5H). However, we did not control for resting membrane potential of the motoneurons and it is possible that differences in driving forces between the motoneuron hid this effect. We observed that the time it takes for the AHP to repolarize to half its amplitude (AHP half-relaxation time [19]), was systematically longer in the S-type motoneurons, followed by FR, followed by FI and FF (Figure 5B,I). Interestingly, none of the electrophysiological properties shown in Figure 5 differed between FI and FF motoneurons (A full comparison of all the properties is available in Supplemental Table 2 and Supplemental Figure 3).

**Figure 5.**
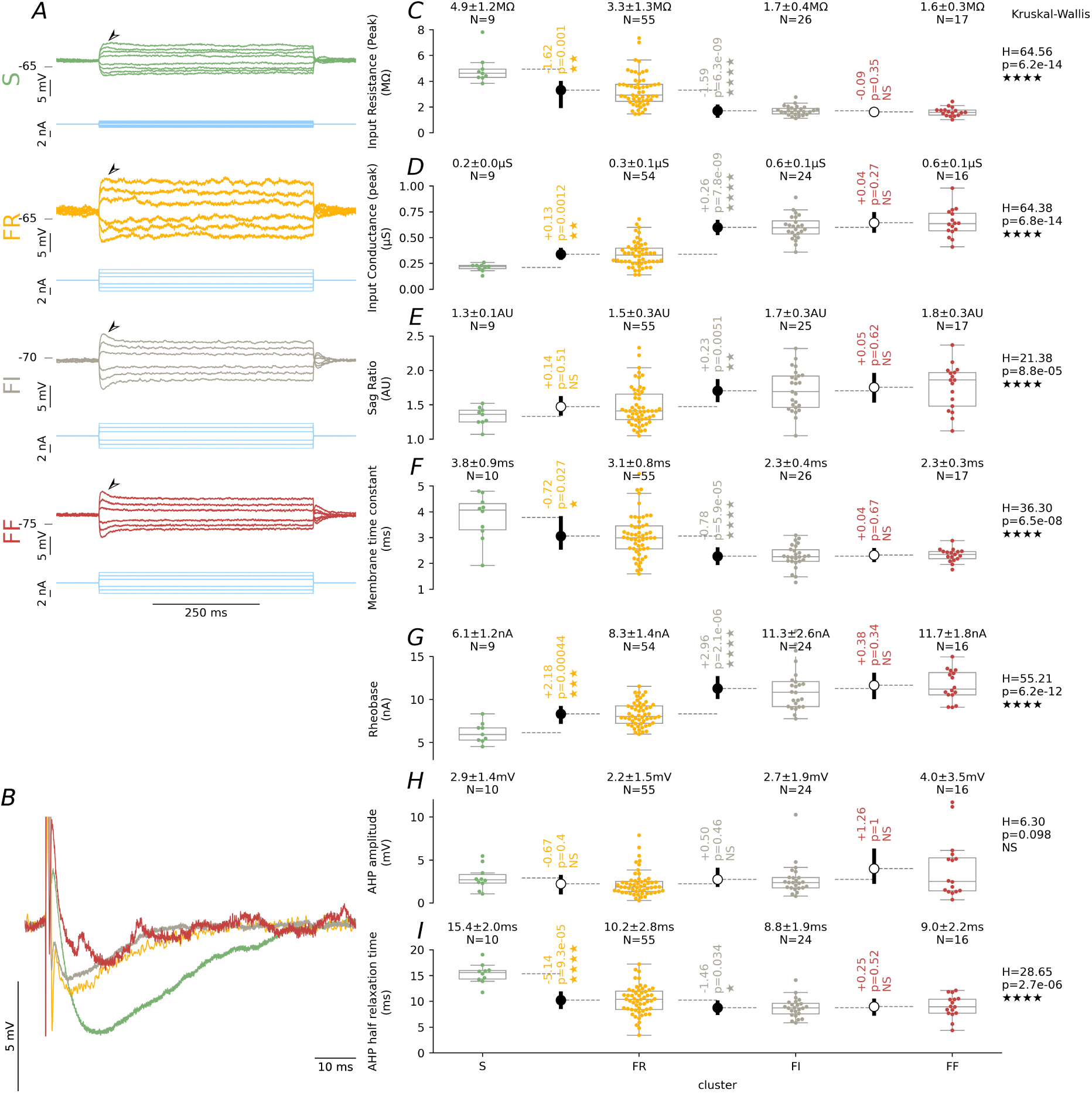
Distribution of the major electrophysiological properties across the different types. **A** Response of four motoneurons (top traces in each sub-panel) to 500 ms square pulses of current (shown below each voltage trace) used to measure the input resistance of the motoneuron. Note the difference in the amount of sag in the response between the different types of motoneurons (arrowheads). Note that the current intensity is not the same in all recordings. **B** Superimposed traces showing the after-hyperpolarization following an intrasomatic spike in 4 representative motoneurons. Note the difference of time course between the S-type AHP and the others. Traces are averages of 5-10 spikes, and were aligned to the baseline potential before the spike. Spikes were truncated for illustration purposes. **C-I** Distribution of the major electrophysiological properties across the different types. For each property (row) and cluster (columns), we show the individual data points superimposed over a boxplot that shows the three quartile values of the distribution along with extreme values. The “whiskers” extend to points that lie within 1.5 x inter-quartile distance of the lower and upper quartile. Above each boxplot, we report the mean ± standard deviation of each of the cluster sub-population. On the far right of the plot is the result of the Kruskal-Wallis H-test for independent samples, showing the H statistic and the *p* value. In between each cluster, we show the difference of the means (shown with dashed lines) between the group to the left and the right, with its associated 95 % confidence interval. The *p* value corresponds to the result of the pairwise Mann-Whitney U Test post-hoc test with Holm adjustment for family-wise error rate. If the Kruskal-Wallis test shows that the four populations have different means, and the Mann-Whitney test is statistically significant, then the difference of means is shown with a filled symbol, otherwise, it is shown with an empty symbol. *p* < 0.05: ★, *p* < 0.01: ★★, *p* < 0.001: ★★★, *p* < 0.0001: ★★★★.

We studied the firing of the motoneurons by injecting triangular ramps of current through the microelectrode (*f-I* curves). As expected from the properties reported above and Henneman’s size principle, S type motoneurons were recruited with a smaller current than FR, which, in turn, were recruited at lower current than the FI and FF motoneurons (Figure 6A-C). Once recruited, mouse motoneurons generally exhibit a “sub-primary range” during which the firing frequency climbs steeply and with a high variability, followed by a more linear “primary range” [19]. We did not detect a difference in the slope of the primary range (also called “gain”) between the different types of motoneurons. The force-frequency curves (Figure 6A bottom row) show that the force increased rapidly as the firing increased and tended to reach a plateau before the top of the ramp, where increased in firing frequency no longer translated into an increase of force. This is consistent with our previous observations, which showed that the majority of the force of a motor unit is recruited during the sub-primary range [32], and is most likely linked to the fact that, in adult mice, the duration of the after-hyperpolarization is shorter than the duration of the muscle twitch (Figure 6D). Note that we did not attempt to push the motoneurons up to their absolute maximum firing frequency as this leads to spike inactivation that is often difficult to recover from. Therefore the maximal frequencies reported here are underestimations of the motoneuron maximal firing frequency.

**Figure 6.**
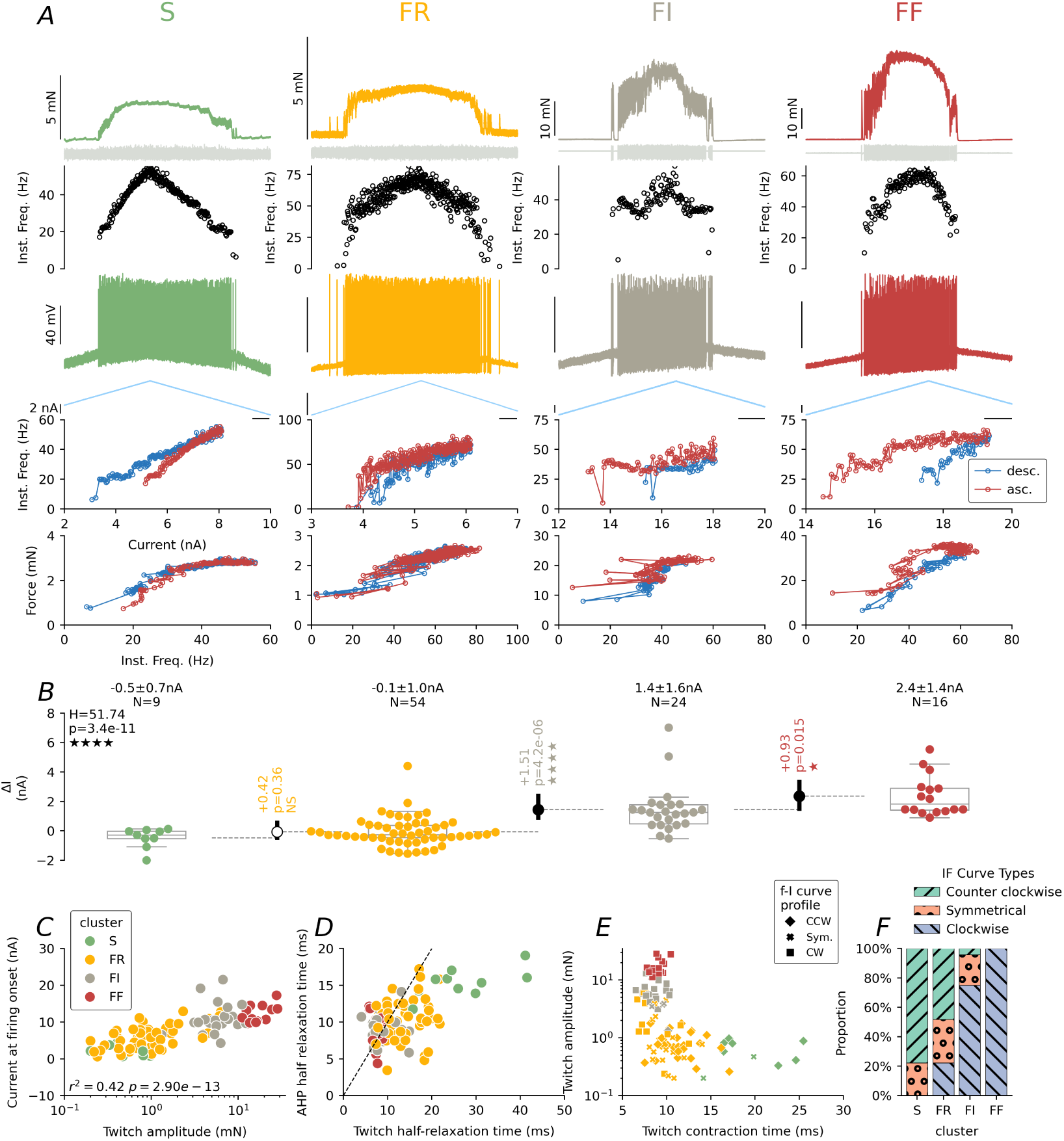
Firing profiles of the motoneurons in each of the four clusters. **A** Representative response of 4 motoneurons to a triangular ramp of current. For each motoneuron, we show (from top to bottom): the force output, the MUAP recording, the instantaneous firing frequency, the voltage trace, the injected current, the corresponding force vs. current curve (*f-I* curve) and the force vs. frequency curve. Note that the scale bars for the force traces are not the same across the recordings. Scale bars for voltage and current are the same in all recordings. Horizontal scale bars: 1 s. **B** Distribution of the value of ΔI across the different types. Same organization as in Figure 5. *p* < 0.05: ★, *p* < 0.01: ★★, *p* < 0.001: ★★★, *p* < 0.0001: ★★★★. **C** Scatter plot of the onset current, i.e. the current required to start firing on the ascending ramp, vs. the twitch amplitude (on a logarithmic scale). **D** Scatter plot of the after-hyperpolarization half-relaxation time (which is proportional to the AHP duration) vs. the twitch half-relaxation time (proportional to the twitch duration). The dashed line represents the identity line. Most data points lie below the identity line, showing the twitch lasts longer than the AHP. **E** Scatter plot showing the distribution of the different profiles of *f-I* curves among the different clusters of motor units. **F** Proportion of the different types of *f-I* curves in the four clusters of motoneurons.

During the descending phase of the ramp, firing either stopped at a de-recruitment current intensity higher than the recruitment current (clockwise hysteresis, ΔI>0), stop at roughly the same current intensity (symmetrical, ΔI∼0), or continue firing until the de-recruitment current is lower than the recruitment current (counter-clockwise hysteresis, ΔI<0). As shown in Figure 5 and Figure 6D-E, there was a systematic difference in firing profile between the different types. The vast majority (∼80 %) of the S-type motoneurons displayed a counter-clockwise hysteresis and a negative ΔI. The remaining 20 % of the *f-I* curves were symmetrical (ΔI∼0). About half of the FR units displayed a counter-clockwise hysteresis, and a small proportion (20 %) of mo-toneurons displaying a clockwise hysteresis. However, this clockwise profile was not distributed uniformly among the FR motoneurons. There was a relationship between the amplitude of the twitch of a motor unit and the value of ΔI (𝑟^2^ = 0.44, p = 2.04⋅10−14, Supplemental Figure 3), indicating that the FR motoneurons displaying a clockwise hysteresis belonged to the largest motor units. Accordingly, the FI and FF units tended to display a majority of clockwise hystereses (Figure 6D-E).

During early post-natal development, motoneurons exhibit two distinct patterns of firing in response to a long pulse of current with an intensity just above rheobase. They either fire immediately at the onset of the pulse, and their firing then adapts over the course of the pulse (*immediate* firing pattern), or they do not fire immediately, but instead, the membrane potential rises slowly during the pulse, until the motoneuron starts to fire with a delay (measured from the pulse onset) and with a firing rate that increases thereafter (*delayed* pattern) [22]. Studies on this behavior suggest that the delayed pattern of firing is caused by the presence of Kv1.2 channels [45] and is observed in large motoneurons, with a high input conductance, and short AHP, and a specific pattern of molecular marker expression, which suggest that they are F-type motoneurons [22]. We tested this behavior in the adult mouse in 25 motoneurons. We found that the vast majority of motoneurons (N=18, 72 %) has a delayed pattern of firing in response to a 5 s-long current pulse at an intensity just above rheobase (Figure 7A2). In fact, we only found two motoneurons that seem to have an immediate firing pattern (8 %). Contrary to the results observed in neonatal animals, motoneurons with a delayed firing pattern were not limited to the fast-type motor units, but four out of the four S-type motoneurons tested exhibited a delayed firing pattern (Figure 7). In the largest motor units (Figure 7B), with the smallest input resistance (Figure 7C), we observed that the motoneurons would fire intermit-tently throughout the 5 s-long pulse, and the firing frequency was very irregular (Figure 7A1, “stuttering” pattern), precluding their classification in either “immediate” or “delayed”. Because of their large size, these motoneurons required higher current intensities to fire, and it is unclear at this stage if this pattern of firing is linked to difficulties in injecting such large currents through the microelectrode, of if it reflects the natural propensity of these units to fire in bursts, rather than tonically [13], [46], [47].

**Figure 7.**
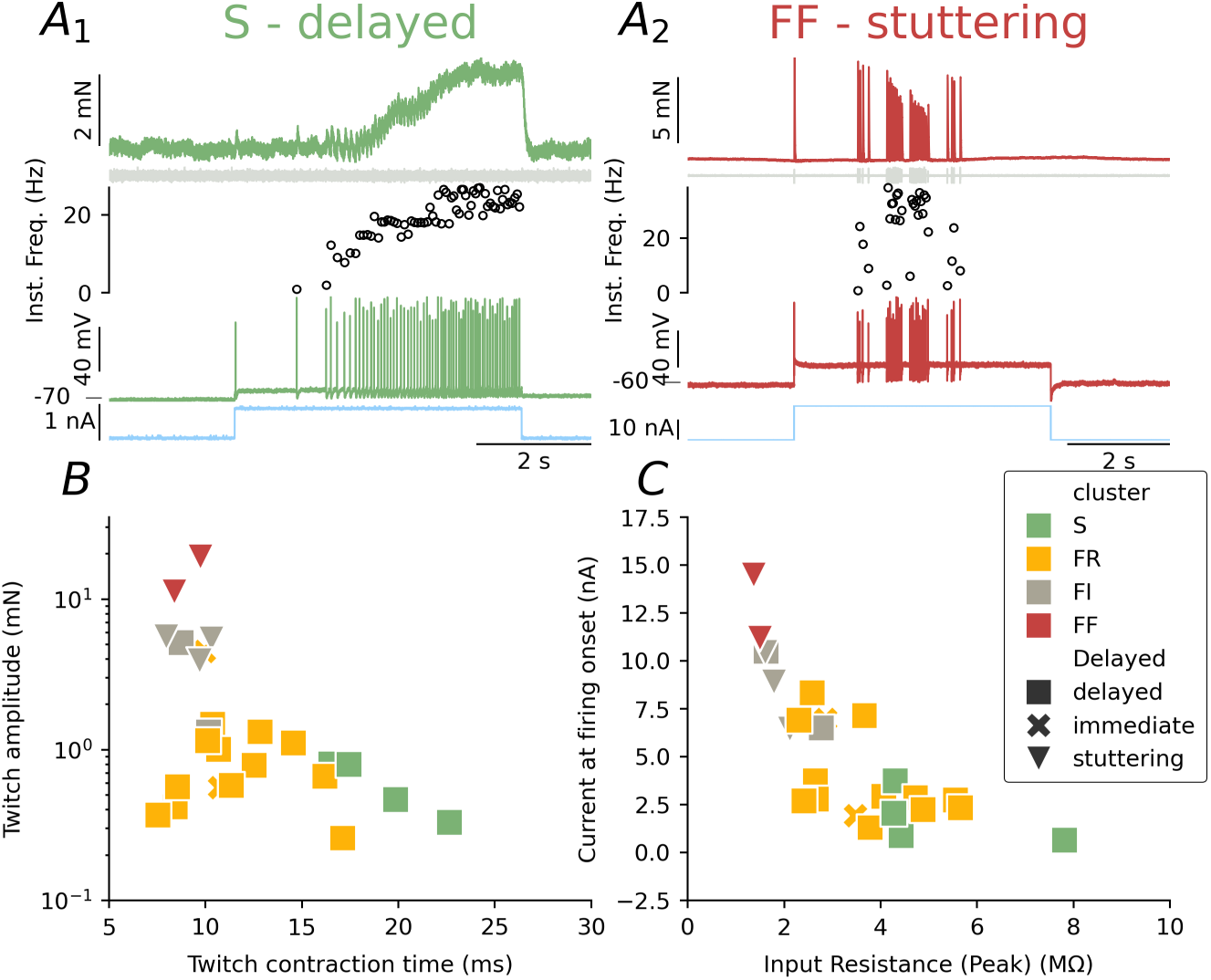
Most motoneurons respond to a long pulse of current with a delayed firing profile. **A** Representative recordings of the response of two motoneurons to a 5 s-long pulse of current at an intensity very close to the rheobase. For each motoneuron, we show (from top to bottom): the force output, the MUAP recording, the instantaneous firing frequency, the voltage trace, and the injected current. **A_1_** Response of a S motoneuron to a 0.8 nA pulse of current. Although it fired a single action potential at the onset of the pulse, its firing stopped for a long period of time before starting again, and the firing frequency increased over the duration of the pulse, characteristic of a *delayed* firing pattern. **A_2_** Response of a FF motoneuron to a 17 nA pulse of current. This neuron was not firing any action potential in response to a 16 nA, but at 17 nA, it was firing in an irregular pattern (“stuttering”) in the middle of the pulse, and it was impossible to assign it to either the immediate or delayed category. **B** Scatter plot of the twitch amplitude vs. the twitch contraction time showing the motoneurons that fired with a delayed pattern with a square, those that fired with an immediate pattern with a cross. There was a third category, termed “stuttering”, which is represented with a downward triangle. The grey dots in the background represent the rest of the population. **C** scatter plot of the onset current, i.e. the current required to elicit firing on the ascending phase of a triangular ramp of current, vs. the input resistance. Same symbols as in B.

### Physiological type prediction

We generated a predictive statistical model that would be capable of predicting the type of a motor unit solely based on the electrophysiological properties of its motor unit. We chose to focus on a set of electrophysiological properties that (1) we believe to be highly relevant to the physiology of the motoneurons, (2) are commonly recorded during our experiments as well as others’, and (3) were featured in a significant proportion of our motoneurons. These features include passive properties (input resistance, time constant, etc.) spike properties (height, width, AHP), and firing properties measured on the response to a triangular ramp of current. Using a total of 23 features, we generated a dataset of 103 motoneurons with no missing data, and a class distribution of nine percent S-type motoneurons (N=9), 52 % FR motoneurons (N=54), 23 % FI (N=24) and 16 % FF (N=16). This distribution was similar to the distribution observed in our full sample (𝜒^2^ (3) = 1.996, p = 0.57).

We performed a systematic search for the model yielding the best predictive power of the motor unit type from motoneuron electrical properties. A multinomial logistic regression classifier was chosen for subsequent analysis as it provided one of the best overall performance, and it is relatively easy to interpret. The model, which included standardized continuous features and one-hot encoded categorical variables, achieved a mean balanced accuracy of approximately 0.75 across five-fold cross-validation, outperforming other tested algorithms such as random forest, gradient boosting, and *k*-nearest-neighbor.

Inspection of the confusion matrix revealed that the model perfectly classified the S motor units and was able to identify FR units fairly well (F1 score = 0.84), indicating that the features captured the distinctive electrophysiological profile of these more excitable cells (high input resistance, low rheobase, and longer afterhyperpolarization durations). In contrast, the classifier struggled to distinguish between FI and FF units (F1 score = 0.56 and 0.55, respectively). As shown above, the electrical properties of these two types were largely overlapping and were not significantly different, which likely explain their higher rate of mutual misclassification.

When the task was simplified to a three-class problem, merging the two largest types (FF and FI) together while retaining FR and S as separate categories, the model’s performance improved substantially. In this situation the classifier achieved a balanced accuracy close to 90 %, with clear separation between low-threshold (S *vs.* FR) and high-threshold (FI+FF) motoneurons. This finding suggests that while the electrical properties alone are insufficient to finely discriminate among the two largest motor unit types (Figure 5), they robustly encode the major functional gradient from slow, fatigue-resistant to fast, fatigable units.

We calculated the odds ratio associated with each feature in order to estimate their contributing to the separation of motoneuron clusters (Figure 8C). For the S cluster, the highest odds ratios were found for the AHP half-relaxation time (OR = 1.32 [95%CI 1.18–1.45]), width of the spike (OR = 1.17 [95%CI 1.04–1.31]), and input resistance (OR = 1.16 [95%CI 1.09–1.24]), whereas features such as maximum frequency (OR = 0.72 [95%CI 0.67–0.77]) and rheobase (OR = 0.83 [95%CI 0.77–0.89]) showed negative associations. These findings indicate that the S cluster were characterized by broader spikes (although we did not detect a statistical difference at the population level Supplemental Table 2), higher input resistances, and lower currents, whereas the cluster for the largest units was associated with shorter afterhyperpolarization and higher firing rates. In the FF+FI cluster, the strongest positive associations were observed for input conductance (OR = 1.25 [95%CI 1.19–1.31]), and the current at firing onset and offset (OR = 1.19 [95%CI 1.14–1.23] and OR = 1.20 [95%CI 1.15–1.25], respectively), indicating that neurons with larger conductances and higher current thresholds were more likely to belong to this group. A complete list of the odd ratios is provided in Supplemental Table 4.

**Figure 8.**
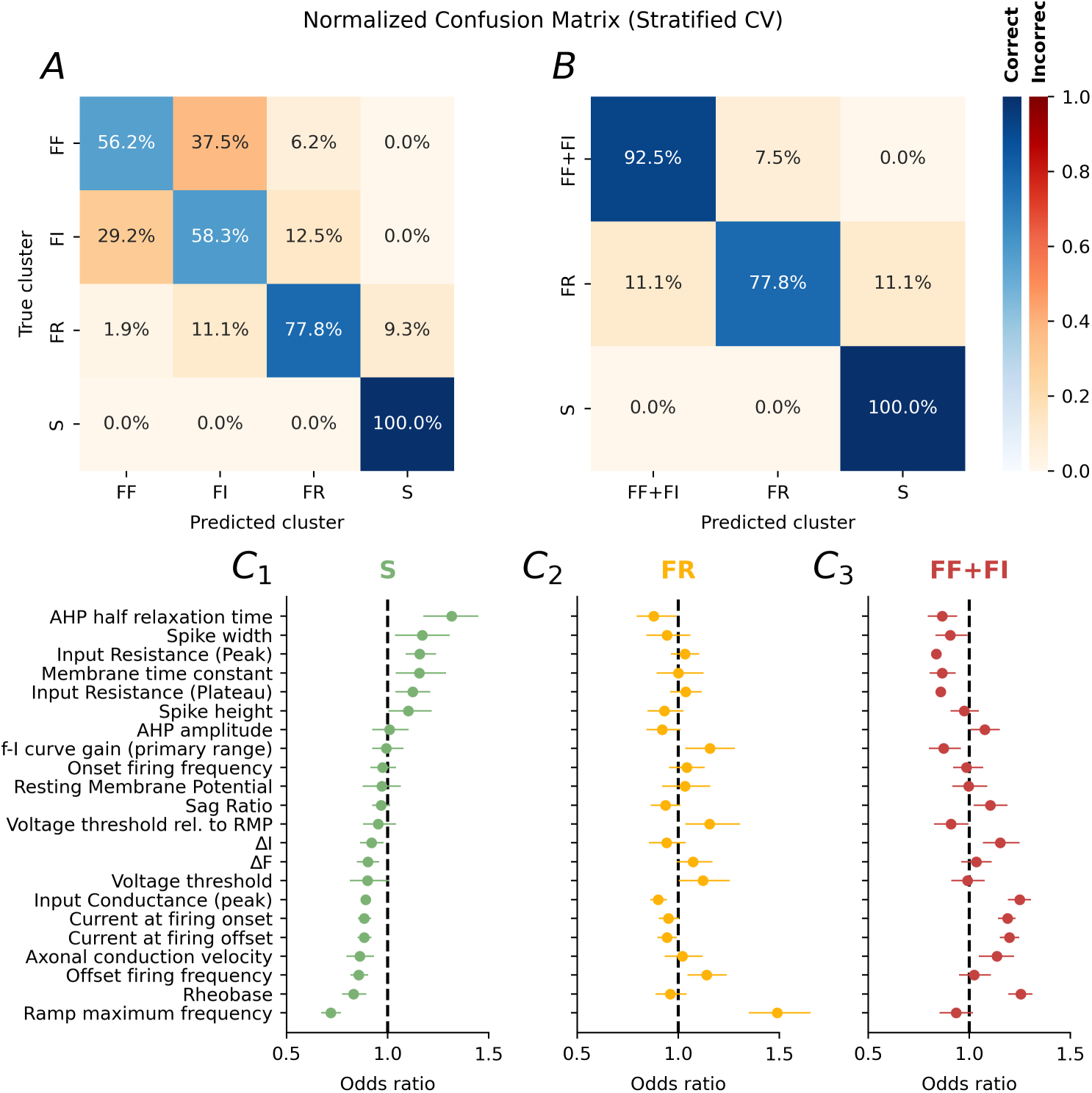
Prediction of motor unit types using machine learning. **A** Normalized confusion matrix for the four-class classifier (FF, FI, FR, S). The main diagonal (blue) shows the proportion of motor units correctly classified, while the off-diagonal squares (red) show the proportion of units that were misclassified. The matrix shows high accuracy for classes FR and S, but large overlap between FF and FI. **B** Confusion matrix for the simplified three-class version (FF/FI vs FR vs S, same organization as in A), demonstrating improved separability and overall accuracy. **C** Odds ratios and 95% confidence intervals for electrophysiological predictors of cluster identity derived from a multinomial logistic regression model. Features were sorted by descending odds ratios in the S cluster. Each point represents the odds ratio for a given electrophysiological property, and horizontal bars indicate the 95% confidence interval estimated by bootstrapping (n = 1000 resamples). Values greater than 1 indicate that higher feature values increase the likelihood of belonging to that cluster, whereas values less than 1 indicate the opposite. Colors denote the motoneuron clusters (S **C_1_**, FR **C_2_**, FF+FI **C_3_**). The dashed vertical line at 1.0 marks the point of no effect.

To improve the generalization power of the model, we sought to reduce the number of input features required while retaining predictive accuracy. Starting from our initial set of 23 electrophysiological parameters, we applied recursive feature elimination with cross-validation (RFECV) to identify the minimal subset of features that preserved model performance. The results indicated that the model could be reduced to 13 features without loss of accuracy, and more notably, that almost as good performance could be achieved with just four features: input conductance, rheobase, AHP half-relaxation time, and maximal frequency. The model was retrained with these four predictors, achieving an overall accuracy of 85.4% (Figure 9). The odds ratios showed that the S cluster identity is strongly dependent on the AHP half-relaxation time (OR = 2.70 [95%CI 1.84–4.20]), while larger input conductances, rheobase or maximum frequency have a negative impact on this cluster Figure 9B1. On the other hand, input conductance was the largest contributor (OR = 4.03 [95%CI 2.72–6.28]) to the large motor unit (FF+FI) group (Figure 9B3). The FR group was associated with high firing frequencies (OR = 3.21 [95%CI 2.26–5.05]) but negatively influenced by high input conductance (OR = 0.57 [95%CI 0.36–0.86])(Figure 9B2). A complete list of the odd ratios is provided in Supplemental Table 4.

**Figure 9.**
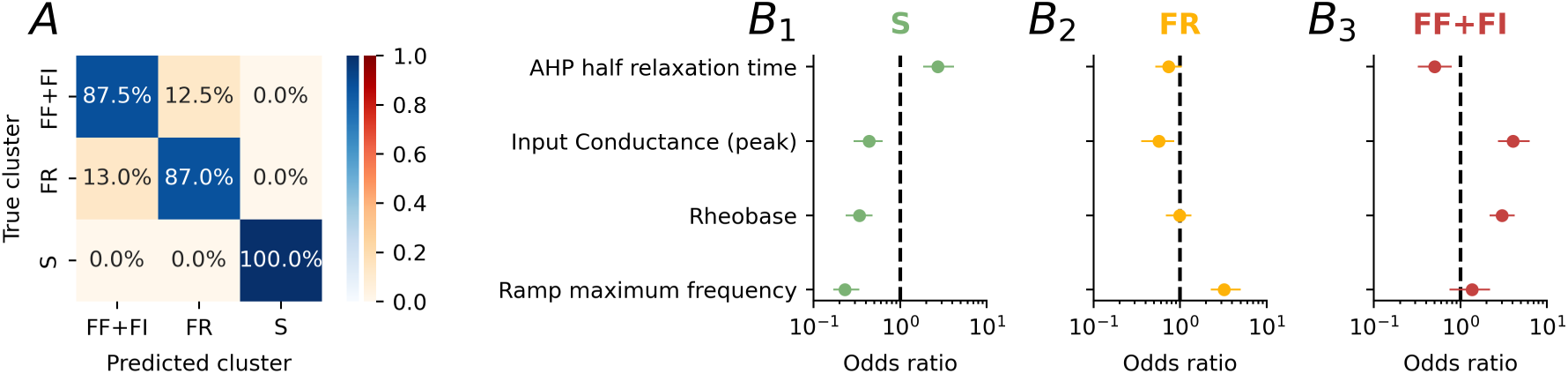
Simplified classifier model. **A** Normalized confusion matrix for the model based on only four electrophysiological features. Same organization as in Figure 8A. **B** Odds ratios and 95% confidence intervals for each of the fours electrophysiological predictors of cluster identity used in the model in A. Same organization as in Figure 8C.

In their seminal work on cat medial gastrocnemius motoneurons, Zengel et al. [12] used linear discriminant analysis using combinations of rheobase, input resistance, membrane time constant, and AHP half-decay time to predict motor-unit type. Their approach achieved remarkably high accuracy (up to 97 %) in distinguishing FF, FR, and S units, particularly when using only rheobase and input resistance. We tested whether their approach could be used on mouse motoneurons. We used their predictors and trained three-class decision-tree models, and validated their performance using a five-fold cross validation approach. We observed lower overall accuracies (≈60–76 % across models, peaking at 0.76 using the product of the input resistance and the membrane time constant) and less separability among classes. While rheobase and input resistance again provided the strongest predictors, no single or combined pair of features reached the discriminatory power of our complete model. The poorer performance of the simpler decision trees likely reflects a more compact and overlapping distribution of motoneuron properties in the mouse compared with the cat [14].

Python code for reproducing the results presented here, as well as our trained model, are freely available. The model can be used with either the full set of 23 properties, or a reduced number of properties (13 or four). The model outputs the most likely class of a motoneuron given the properties provided.

## Discussion

In this article, we provide, for the first time in adult mice, a complete examination of the relationship between the electrical properties of motoneurons and the contractile behavior of their motor units in the mouse Triceps Surae. Force recordings followed by unsupervised clustering identified four physiological groups consistent with the classic slow (S), fast fatigue-resistant (FR), intermediate (FI), and fast fatigable (FF) types. Motoneurons belonging to these clusters differed in rheobase, input resistance, afterhyperpolarization duration, and firing adaptation, demonstrating a tight coupling between electrical and mechanical properties. Machine learning models trained on motoneuron features predicted contractile identity with good accuracy for S and FR types and lower accuracy for FI and FF. These results demonstrate a graded but systematic relationship between motoneuron excitability and muscle contractile phenotype.

The electrophysiological properties of mouse motoneurons observed here are consistent with long-standing expectations derived from Henneman’s size principle [48]. Motoneurons with higher input resistance and longer after-hyperpolarizations exhibited lower rheobase and were associated with slower, more fatigue-resistant motor units, while those with lower resistance and shorter AHPs were linked to faster, more fatigable units. This orderly relationship between excitability and motor unit contractile properties supports the notion of smooth recruitment, in which force is gradually modulated as progressively larger motoneurons are recruited. Our data show that, in mice, the motor unit properties also co-evolve in a continuous manner: as rheobase increased, twitch contraction time decreased and fatigue resistance declined. Principal component analysis placed the motor units along a slow-to-fast continuum rather than into separate clusters with clear boundaries. A continuous organization likely facilitates smooth gradation of force output.

Unexpectedly, the vast majority of motoneurons in our adult in vivo recordings displayed delayed firing in response to just-suprathreshold current injection. This stands in sharp contrast to results from neonatal *ex vivo* spinal cord preparations, where delayed firing was typically restricted to large, fast-type motoneurons associated with high conductance and expression of Kv1.2 channels [22], [45]. The prevalence of delayed firing across motoneuron types suggests differences in ionic channel maturation, temperature, neuromodulatory drive, or recording configuration could each contribute to the discrepancy, highlighting the importance of studying adult motoneurons within their intact physiological context.

### Discrete Motor Unit Types versus a Functional Continuum

The physiological classification of motor units has traditionally relied on grouping them into discrete types. Early work in cats and rats established three or four categories of motor units based on twitch contraction time, maximal force, and fatigue index [3], [12]. This framework has proven useful for describing recruitment order and functional specialization. Yet, both historical and recent studies investigating individual motor unit contractile properties have repeatedly demonstrated that these properties are continuously distributed rather than discretely classified. For instance, Bigland-Ritchie, Fuglevand, and Thomas reported that human thenar motor units display a continuous spectrum of force output and contraction speed, with no clear separation into distinct types [49]. Similarly, studies involving intraneural motor axon stimulation of single motor units, such as those conducted on human toe extensors and in cat distal forelimbs, have established that the variability in contractile characteristics follows a continuous distribution [4], [49], [50], [51], [52]. Motoneuron electrical features such as input resistance, AHP duration, and rheobase vary smoothly along the same axis, producing an overlapping gradient from the smallest, most excitable neurons to the largest, least excitable ones [12], [13], [14], [35], [44]. Although motor units are thought to be constitutes of muscle fibers with homogenous properties, there is also considerable variability in the histochemical profiles of muscle fibers [53], [54], [55], [56], [57], [58], which will similarly contribute to a continuous profile of force recruitment.

While a continuum may better reflect physiology reality, grouping motor units retains practical value. Discrete categories simplify comparison across muscles, species, and experimental conditions. The physiological ranges of contraction time or rheobase differ between muscles such as the Soleus and Extensor Digitorum Longus, making direct comparison difficult without standard labels [5], [59]. Classification into sub-groups provides a shared vocabulary that allows consistent reporting of trends. It also supports statistical analyses when sample sizes are limited. Grouping can highlight shifts in population composition, such as the loss of fast fatigable units during disease progression or training-induced increases in fatigue-resistant units.

### Relevance for studying motor units in diseases

Selective vulnerability is a defining feature of many neuromuscular disorders [16]. Fast fatigable motoneurons degenerate earliest in amyotrophic lateral sclerosis (ALS) [60], [61], [62], spinal muscular atrophy [63], Kennedy’s disease [64], and normal aging [65], [66]. Slow and fatigue-resistant units, by contrast, persist until late disease stages.

Traditional analyses often treat motoneurons as a homogeneous pool, averaging properties across all recorded cells. This approach obscures early, subtype-specific changes. By defining electrophysiological boundaries that correspond to functionally distinct units, researchers can examine how disease processes shift each subgroup’s properties. For example, a change in input conductance and resting membrane potential within the FF cluster might indicate selective stress on the largest motoneurons even before overt degeneration occurs [15]. The ability to classify cells based on electrical features alone also makes it possible to identify vulnerable subtypes in preparations where contractile measurements are not feasible, such as *ex vivo* spinal cord preparations [67], [68], [69] or when the use of curare is desirable to improve the stability of the recordings *in vivo*.

### Limitations of the study

One limitation of the present study is that the number of recorded motor units was modest, and some clusters contained fewer observations. This limitation reflects experimental constraints and the natural distribution of motor unit types within the mouse Triceps Surae. Histological and electrophysiological studies indicate that the Gastrocnemius muscles are dominated by fast units, while the Soleus contains more slow and fatigue-resistant fibers [41], [54]. Although numbers were not available for the whole triceps surae, we used counts of muscle fibers [70], [71] and proportions of muscle fiber types [55], [72] in the soleus and gastrocnemius muscles to estimate that the distribution over the whole Triceps Surae muscle is 5% type I, 14% type IIa, 6% type IIx, and 75% type IIb. The relative scarcity of slow units in our sample is therefore expected rather than a sampling artifact, and we actually recorded a larger proportion of S-type motoneurons than expected from looking at the proportion of fiber types. This discrepancy can be explained by the fact that the innervation ratios (i.e. number of muscle fibers innervated by each motoneuron) is larger in FF and FR units than in S units [73], [74], [75].

Another limitation is that the model was derived from only one muscle group (Triceps Surae). That muscle is highly physiologically relevant for gait, balance, motor control and clinical research in animal models and humans, and has the advantage to contain all types of muscle fibers in substantial proportions, whereas other muscles may be devoid of one type or another [15]. Nevertheless, electrical properties depend on motoneuron size and muscle function, which differ between proximal and distal muscles. Although the size principle and the overall continuum of excitability appear conserved across muscles, quantitative relationships may vary. Applying the same model to other muscles will test its generality and reveal whether distinct muscles share a common electrophysiological-to-contractile mapping.

## Conclusion

Despite these limitations, the present study is the first to integrate contractile, electrophysiological, and computational analyses of identified mouse motor units within the same preparation. These results have both physiological and translational relevance, establishing a foundation for future work on motor unit diversity, recruitment, and degeneration. This framework opens the way for systematic studies of motor unit function in health and disease using unified, quantitative criteria.

### Data availability

All analysis scripts, data-preprocessing pipelines, and trained classifier models are available at this repository.

## Acknowledgments

This work was supported by NIH NINDS R01NS110953 (MM). The authors thank Daniel Zytnicki and Guillaume Caron for helpful discussions on the data and their comments on the manuscript.

## Author contribution

MdLMS: Investigation, Data analysis, Writing - Review & Editing; RMA: Methodology, Formal analysis, Writing - Review & Editing; EJR: Investigation, Data analysis, Writing - Review & Editing; RDIM: Resources, Writing - Review & Editing; NK: Methodology, Formal analysis, Supervision, Funding acquisition; MM: Conceptualization, Methodology, Software, Investigation, Data Curation, Data analysis, Visualization, Writing - Original Draft, Supervision, Funding acquisition;

## Declaration of interests

The authors declare no competing interests.

**Supplemental Table 1** – Table of the contractile properties in each of the four clusters of motor units. In each cell, we report the mean value +/- standard deviation as well as the range of the values and the number of observations. Then, we show the result of the Kruskal-Wallis H-test for independent samples, showing the H statistic and the *p* value. Then, for each pair of cluster, we show the effect size of the difference (Hedge’s *g* with 95% confidence interval) and the result of the pairwise Mann-Whitney U Test post-hoc test with Holm adjustment for family-wise error rate for each pair of cluster. *p* < 0.05: ★, *p* < 0.01: ★★, *p* < 0.001: ★★★, *p* < 0.0001: ★★★★.

**Supplemental Table 2** – Table of the electrophysiological properties in each of the four clusters of motor units. In each cell, we report the mean value +/- standard deviation as well as the range of the values and the number of observations. Then, we show the result of the Kruskal-Wallis H-test for independent samples, showing the H statistic and the *p* value. Then, for each pair of cluster, we show the effect size of the difference (Hedge’s *g* with 95% confidence interval) and the result of the pairwise Mann-Whitney U Test post-hoc test with Holm adjustment for family-wise error rate for each pair of cluster. *p* < 0.05: ★, *p* < 0.01: ★★, *p* < 0.001: ★★★, *p* < 0.0001: ★★★★.

**Supplemental Figure 2** – Scatter plots of the various contractile properties (rows) as a function of the twitch amplitude, twitch contraction time, and fatigue index (columns). Color code corresponds to the different clusters of motor units

**Supplemental Figure 3** – Scatter plots of the various electrophysiological properties (rows) as a function of the twitch amplitude, twitch contraction time, and fatigue index (columns). Color code corresponds to the different clusters of motor units

**Supplemental Table 3** – Table of the Principal Components loadings for each of the first 5 principal components.

**Supplemental Table 4** – Table showing the Odds Ratio values for the complete model, and the reduced model with 4 input features.

## Notes

https://zenodo.org/records/17632990

### Competing Interest Statement

The authors have declared no competing interest.

