## Supplementary figures and images for "Machine-Learning Classification of Motor Unit Types in the Adult Mouse"

### Supplemental Figure 1

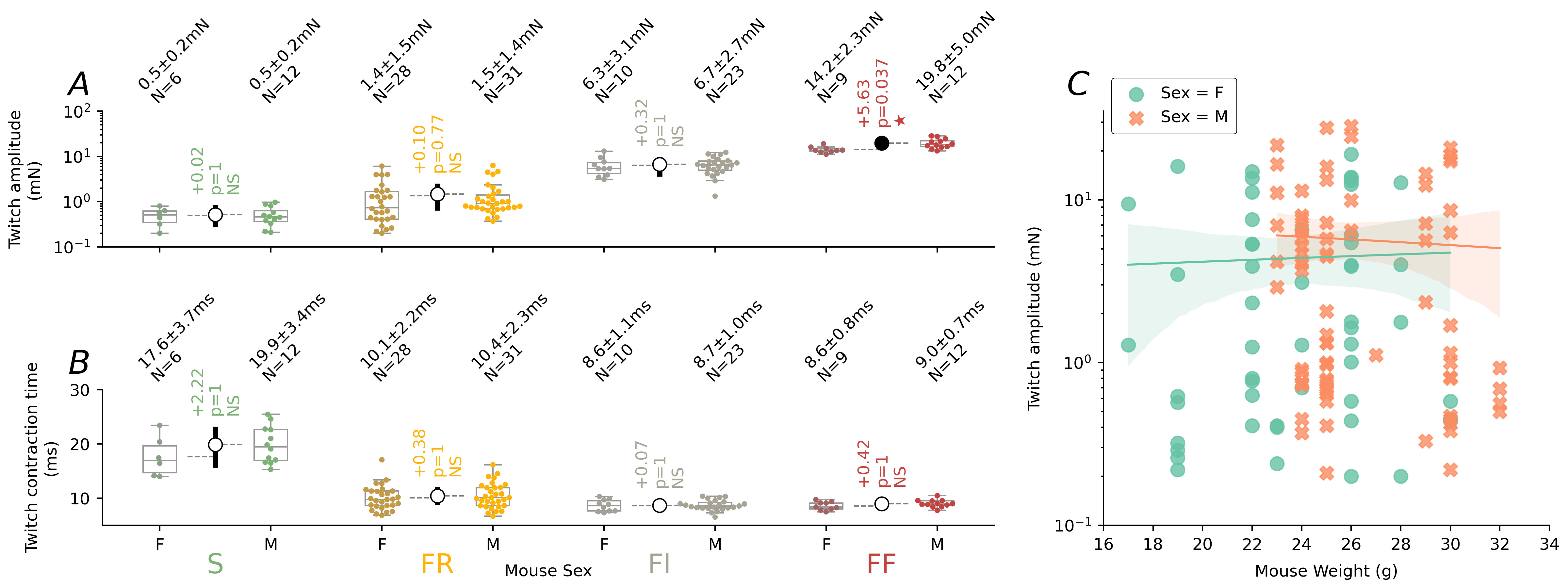
